# Optimal foraging and the information theory of gambling

**DOI:** 10.1101/497198

**Authors:** Roland J. Baddeley, Nigel R. Franks, Edmund R. Hunt

**Affiliations:** School of Experimental Psychology, University of Bristol, 12a Priory Road, Bristol, BS8 1TU, UK; School of Biological Sciences, University of Bristol, Life Sciences Building, 24 Tyndall Avenue, Bristol, BS8 1TQ, UK; School of Computer Science, Electrical and Electronic Engineering, and Engineering Mathematics, Merchant Venturers Building, 75 Woodland Road, Bristol, BS8 1UB, UK

**Keywords:** Movement ecology, collective behaviour, Bayesian methods, Markov chain Monte Carlo, Lévy foraging, sociobiology

## Abstract

At a macroscopic level, part of the ant colony life-cycle is simple: a colony collects resources; these resources are converted into more ants, and these ants in turn collect more resources. Because more ants collect more resources, this is a multiplicative process, and the expected logarithm of the amount of resources determines how successful the colony will be in the long run. Over 60 years ago, Kelly showed, using information theoretic techniques, that the rate of growth of resources for such a situation is optimised by a strategy of betting in proportion to the probability of payoff. Thus, in the case of ants the fraction of the colony foraging at a given location should be proportional to the probability that resources will be found there, a result widely applied in the mathematics of gambling. This theoretical optimum generates predictions for which collective ant movement strategies might have evolved. Here, we show how colony level optimal foraging behaviour can be achieved by mapping movement to Markov chain Monte Carlo methods, specifically Hamiltonian Markov chain Monte Carlo (HMC). This can be done by the ants following a (noisy) local measurement of the (logarithm of) the resource probability gradient (possibly supplemented with momentum, i.e. a propensity to move in the same direction). This maps the problem of foraging (via the information theory of gambling, stochastic dynamics and techniques employed within Bayesian statistics to efficiently sample from probability distributions) to simple models of ant foraging behaviour. This identification has broad applicability, facilitates the application of information theory approaches to understanding movement ecology, and unifies insights from existing biomechanical, cognitive, random and optimality movement paradigms. At the cost of requiring ants to obtain (noisy) resource gradient information, we show that this model is both efficient, and matches a number of characteristics of real ant exploration.

## Introduction

Life has undergone a number of major organisational transitions, from simple self-replicating molecules into complex societies of organisms (Maynard Smith and Szathmary, 1995). Social insects such as ants, with a reproductive division of labour between the egg-laying queen and non-reproductive workers whose genetic survival rests on her success, exemplify the highest degree of social behaviour in the animal kingdom: ‘true’ sociality or eusociality. The workers’ cooperative genius is observed in diverse ways (Camazine et al., 2001) from nest engineering (Dangerfield et al., 1998) and nest finding (von Frisch, 1967), to coordinated foraging swarms (Franks, 1989) and dynamically adjusting living bridges (Reid et al., 2015). This has inspired a number of technological applications from logistics to numerical optimisation (Dorigo and Gambardella, 1997; Karaboga and Basturk, 2007). All of these behaviours may be understood as solving particular problems of information acquisition, storage and collective processing in an unpredictable and potentially dangerous world (Detrain et al., 1999). Movement (the change of the spatial location of whole organisms in time) is intrinsic to the process. Here we consider how optimal information processing is mapped to movement, at the emergent biological levels of the organism and the colony, the ‘superorganism’. We develop a Bayesian framework to describe and explain the movement behaviour of ants in probabilistic, informational terms, in relation to the problem they are having to solve: the optimal acquisition of resources in an uncertain environment, to maximise the colony’s geometric mean fitness (Orr, 2009). The movement models are compared to real movement trajectories from *Temnothorax albipennis* ants.

### Operationalizing conceptual animal movement frameworks

Scientists have studied animal movement for many years from various perspectives, and in recent years attempts have been made to unify insights into overarching frameworks. One such framework has been proposed by Nathan et al. (Nathan et al., 2008). We describe it briefly to set the research context for the reader. Their framework identifies four components in a full description: the organism’s internal state; motion capacity; navigational ability; and influential external environmental factors. This framework also characterises existing research as belonging to different paradigms, namely ‘random’ (classes of mathematical model related to the random walk or Brownian motion); ‘optimality’ (relative efficiency of strategies for maximising some fitness currency); ‘biomechanical’ (the ‘machinery’ of motion); and the ‘cognitive’ paradigm (how individuals’ brains sense and respond to navigational information). However, scientists have yet to create a theoretical framework which convincingly unifies these components. Frameworks such as Nathan et al. are also focused on the individual and so for group-living organisms, especially for eusocial ones, they are incomplete. The concepts of search and uncertainty also need to be better integrated within foraging theory so that the efficiency of different movement strategies can be evaluated (Giuggioli and Bartumeus, 2010).

Here, we contend that animal foraging (movement) models should be developed with reference to the particular information processing challenges faced by the animal in its ecological niche, with information in this context referring to the realised distribution of fitness-relevant resources: in particular the location and quality of foraging patches, which are unknown *a priori* to the organism(s). Furthermore, an important ‘module’ in any comprehensive paradigm for animal movement is the role of the group and its goals in determining individual movement trajectories; there has been much research on collective behaviour in recent years, with information flow between individuals identified as an important focus of research (Sumpter, 2006). Eusocial insects like ants exhibit a highly advanced form of sociality, even being described as a ‘superorganism’, that is, many separate organisms working together as one (Hölldobler and Wilson, 2009). Their tremendous information processing capabilities are seen clearly in their ability to explore and exploit collectively their environment’s resources. Ants thrive in numerous ecological niches, and alone account for 15–20% of the terrestrial animal biomass on average, and up to 25% in tropical regions (Schultz, 2000).

The collective behaviour of tight-knit groups of animals like ants has been described as collective cognition (Couzin, 2009). Because a Bayesian framework seems natural for a single animal’s decision-making (McNamara et al., 2006), an obvious challenge would seem to be applying its methods to describe the functioning of a superorganism’s behaviour. First, we identify a simple model that describes the foraging problem that ants, and presumably other collectives of highly related organisms, have evolved to solve.

### Placing bets: choosing where to forage

Evolution by natural selection should produce organisms that can be expected to have an efficient foraging strategy in their typical ecological context. In the case of an ant colony, although it consists of many separate individuals, each worker does not consume the food it collects and is not independent, but there is rather a colony-level foraging strategy enacted without central control that ultimately seeks to maximise colony fitness (Giraldeau and Caraco, 2000). Following the colony founding stage comes the ‘ergonomic’ stage of a colony’s life cycle (Oster and Wilson, 1978). This is when the queen is devoted exclusively to egg-laying, while workers take over all other work, including collecting food. Thus the colony becomes a ‘growth machine’ (Oster and Wilson, 1978), whereby workers collect food to increase the reproductive rate of the queen, who transforms collected food into increased biomass or more numerous gene copies. Ultimately, the success or failure of this stage determines the outcome of the reproductive stage, where accumulated ‘wealth’ (biomass) correlates with more offspring colonies. This natural phenomenon has parallels with betting, where the winnings on a game may be reinvested to make a bigger bet on the next game. In the context of information theory, John Kelly made a connection between the rate of transmission of information over a communications channel, which might be said to noisily transmit the outcome of a game to a gambler while bets can still be made, and the theoretical maximum exponential growth rate of the gambler’s capital making use of that information (Kelly, 1956). To maximise the gambler’s wealth over multiple (infinite) repeated games, it is optimal to bet only a fraction of the available capital each turn across each outcome, because although betting the whole capital on the particular outcome with the maximum expected return is tempting, any losses would quickly compound over multiple games and erode the gambler’s wealth to zero. Instead, maximising logarithmic wealth is optimal, since this is additive in multiplicative games and prevents overbetting. Solving for this maximisation results in a probability matching or ‘Kelly’ strategy, where bets are made in proportion to the probability of the outcome (Cover and Thomas, 2006). For instance, in a game with two outcomes, one of 20% probability and one of 80% probability, a gambler ought to bet 20% of his wealth on the former and 80% on the latter. This does not depend on the payoffs being fair with respect to the probabilities of the outcome, or 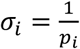, which in the aforementioned case would be 5 and 1.25. Instead it simply requires fair odds with respect to some distribution, or 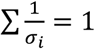 where *σ*_*i*_ is the payoff for a bet of 1, so they could for instance be 2 to 1 or uniform odds in the case of a game with two outcomes (see supplementary Methods). For the purposes of our foraging model, we can simply impose the constraint of fair odds, and any distribution of real-life resource payoffs can be mapped to this when renormalized.

In the case of ants choosing where to forage, the probability matching strategy can be directly mapped onto their collective behaviour. With two available foraging patches having a 20% and 80% probability of food being present at any one time, the superorganism should match this probability by deploying 20% and 80% of foragers to the two sites (though it is also possible to follow a Kelly strategy while holding back a proportion of wealth; see supplementary methods). Regardless of the particular payoff *σ*_*i*_ available at each site, provided 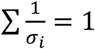 this strategy is optimal over the long term, with the evolutionary time scale of millions of years favouring its selection. Figure 1 shows a simulated comparison of the Kelly strategy where probabilities of receiving a resource payoff are matched, regardless of the payoff, with a strategy that allocates foragers proportional to the one-step expected return *p*_*i*_*σ*_*i*_, which does take the payoff into account.

**Figure 1.**
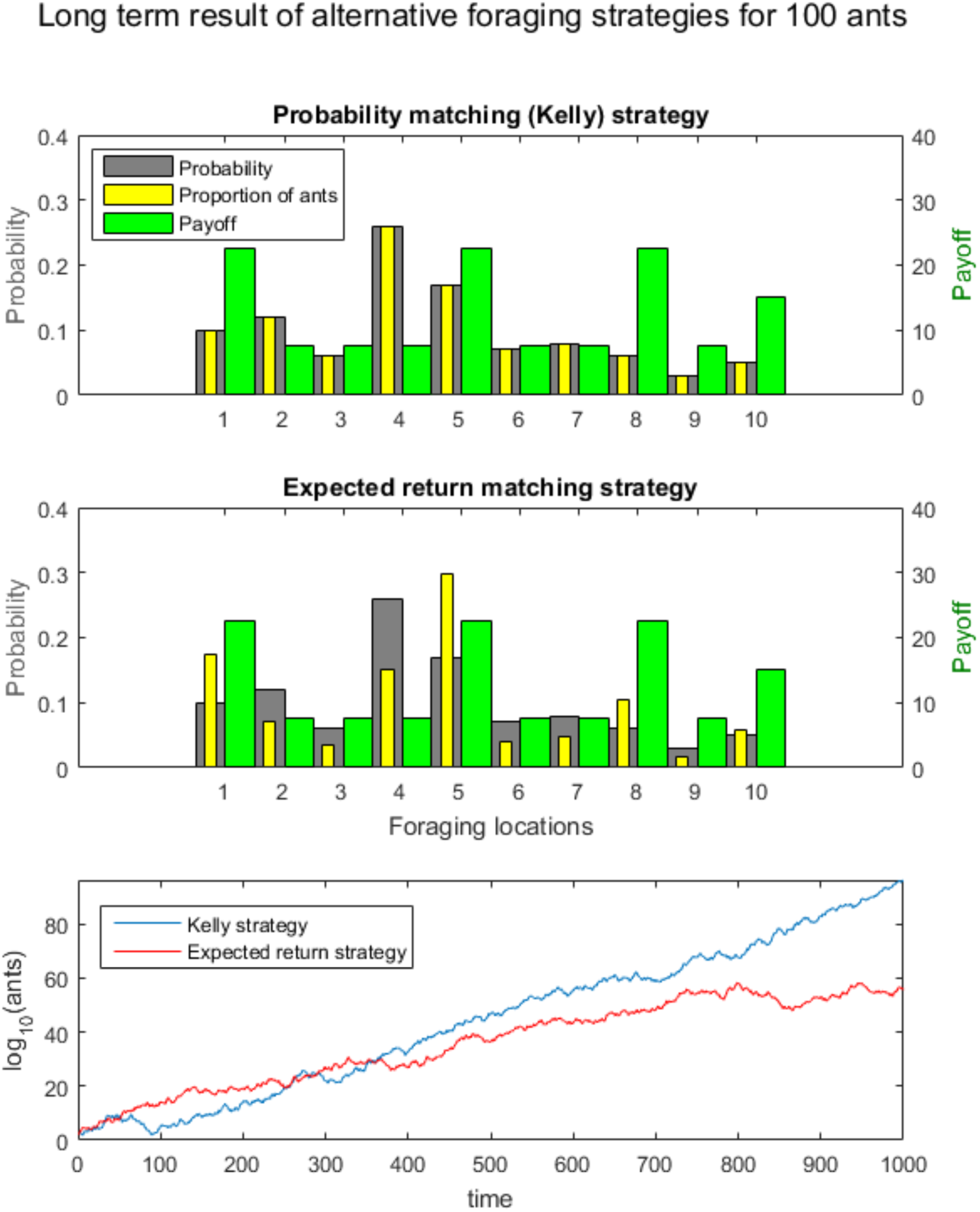
A comparison of the Kelly strategy with an expected return matching strategy, over the long term (identical one-step payoffs for a ‘win’ in both cases). In the top pane (probability matching) the proportion of ants ‘bet’ (yellow bars) matches the probability of success (grey). In the middle pane, the proportion of ants is allocated by the expected return (probability × payoff). The Kelly strategy increasingly outperforms any other strategy as time goes by (bottom pane, example simulation).

Previous analysis of the behaviour of Bayesian foragers versus those modelled using the marginal value theorem indicated that, rather than abandoning a patch when instantaneous food intake rate equals foraging costs, a forager should consider the potential future value of a patch before moving on, even when the current return is poor (Olsson et al., 2006). The priority of resource reliability over immediate payoff in our model, when long-term biomass maximisation is the goal, is itself an interesting finding about superorganismal behaviour; but here we go further and specify models of movement to operationalize this strategy.

Certain methodologies designed to sample from probability distributions – Markov chain Monte Carlo (MCMC) methods – may be used as models of movement that also achieve a probability matching (Kelly) strategy. Exploring the environment and sampling from complex probability distributions can be understood as equivalent problems. MCMC methods aim to build a Markov chain of samples that draw from each region of probability space in correct proportion to its density. A well-mixed Markov chain is analogous to a probability matching strategy. Once the Markov chain has converged on its equilibrium distribution (the target probability distribution, or resource quality distribution in our ant model) it spends time in each location proportional to the quality or value (probability) of each point.

### A colony-level strategy

There is a central ‘social’ (colony-level) element in attempting to enact a Kelly strategy of allocating ‘bets’ in proportion to the probability of their payoff. This is because it requires a ‘bank’ (collection of individuals) that can be allocated. This logic does not seem to apply when one is thinking of a single individual, which might instead prefer (or need) to pursue high expected returns to survive in the short-term. Therefore, our model is relevant to groups of individuals who have aligned interests in terms of their fitness function – this is notably true in the social insects such as the ants, because workers are (unusually) highly related, or in clonal bacteria, for instance.

However, using MCMC as a model of movement does not, in itself, imply social interactions are necessary. Multiple MCMC ‘walkers’ can sample in parallel from a space and still achieve sampling (foraging patch visitation) in proportion to probability. Nevertheless, social interactions could be highly advantageous in expediting an efficient sampling of the space, through for example ‘tandem running’ (Franks & Richardson, 2006) to sample important areas (Hunt et al., in prep, 2018b), or pheromone trails to mark unprofitable territory (Hunt et al., in prep, 2018c).

### Ant trajectory data

We use our data (Hunt et al., 2016a) from previous work examining the movement of lone *Temnothorax albipennis* ants in an empty arena outside their colony’s nest (Hunt et al., 2016b). *T. albipennis* ants have been used as a model social system for study in the laboratory, because information flow between the environment and colony members, and among colony members, can be rigorously studied. The ants typically have one queen and up to 200-400 workers (Franks et al., 2006). The colony inhabits fragile rock crevices and finds and moves into a new nest when its nest is damaged. With workers being only about 2mm long, relatively unconstrained trajectories of individuals can be tracked on the laboratory workbench (for example, Hunt et al. 2016b). Behavioural state-based models have been developed that account for the flow of individuals between states with differential equations (Sumpter and Pratt, 2003; Pratt et al., 2005), but these lack an account of the ants’ movement processes.

## Methods

We run simulations of our Markov chain Monte Carlo movement models in MATLAB 2015b (pseudocode is available in the supplementary material). Each new model is introduced to explain an important additional aspect of the ants’ empirical movement behaviour.

In our movement data (Hunt et al., 2016a) there are two experimental regimes, one in which the foraging arena was entirely novel to exploring ants, and one in which previous traces of the ants’ activities remained. We use the data from the former treatment, where each ant encounters a cleaned arena absent of any pheromones or cues from previous exploring ants. We restrict our analysis to the first minute of exploration, well before any of the ants have an opportunity to reach the boundary of the arena. Log-binned root mean square displacement is calculated and a linear regression made against log time. A gradient equal to a half indicates a standard diffusion process (Brownian motion) whereas greater than a half indicates superdiffusive movements. This approach to characterising ant search behaviour has been taken in e.g. Franks et al. (2010).

A supplementary Methods section is at the end of the paper.

## Results

We present simulation results from three different models of ant movement. Each model is directly based on a known Markov chain Monte Carlo method (MCMC). This follows the recognition that we can consider the problem of sampling from probability distributions of two continuous dimensions as analogous between animal movement and statistics (for example). The trajectories produced by each model are compared to real ant movement data. The development of MCMC methods from the 1950s onward, to become more efficient, might be considered to parallel the evolutionary history of animal foraging strategies. Some more details on the methods are found in the supplementary Methods section.

### Basic model: Metropolis-Hastings

The first MCMC method to be developed was the Metropolis-Hastings (M-H) algorithm (Metropolis et al., 1953; Hastings, 1970), which is straightforward to implement and still commonly used today.

We are trying to sample from the target probability distribution (resource quality distribution) *P*(*x*) which can be evaluated (observed) for any *x*, at least to within a multiplicative constant. This means we can evaluate a function *P*\*(*x*) such that *P*(*x*) = *P*\*(*x*)/*Z*. There are two challenges that make it difficult to generate representative samples from *P*(*x*). The first challenge is that we do not know the normalising constant *Z* = *∫ d*^*N*^*xP*\*(*x*), and the second is that there is no straightforward way to draw samples from *P* without enumerating most or all of the possible states. Correct samples will tend to come from locations in *x*-space where *P*(*x*) is large, but unless we evaluate *P*(*x*) at all locations we cannot know these in advance (Mackay, 2003).

The M-H method makes use of a proposal density *Q* (which depends on the current state *x*) to create a new proposal state to potentially sample from. Q can be simply a uniform distribution: in a discretized environment these can be *x*^(*t*)^ + [−1,0,1] with equal probability. After a given proposed movement is generated, the animal compares the resource quality at this new location with the resource quality at the previous location. If the new location is superior, it stays in its new location. In contrast, if the resource quality is worse, it randomly ‘accepts’ this new location, or ‘rejects’ this location based on a very simple formula based on the ratio of resource quality (if it is far worse, the animal very rarely fails to return, whereas if it is not much worse, it often accepts this mildly inferior location – see also supplementary methods). What is important about this extremely simple algorithm is that, as long as the environment is ergodic (all locations can potentially be reached), given time, the exploring animal will visit each location eventually. Visits will be made with a probability proportional to its resource quality: it will execute an optimal Kelly exploration strategy. The problem here, however, is the time taken. Whilst the M-H method is widely used for sampling from high-dimensional problems, it has a major disadvantage in that it explores the probability distribution by a random walk, and this can take many steps to move through the space, according to 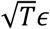 where T is the number of steps and *ϵ* is the step length. *T. albipennis* ants were found to be engaged in a superdiffusive search in an empty arena (supplementary Methods), and similarly MCMC methods also have been developed to explore probability space more efficiently.

### Introducing momentum: Hamiltonian Monte Carlo (HMC)

Random walk behaviour is not ideal when trying to sample from probability distributions, since it is more time-consuming than necessary. One popular method for avoiding the random walk-like exploration of state space is Hybrid Monte Carlo (Duane et al., 1987), also known as Hamiltonian Monte Carlo (HMC). This simulates physical dynamics to preferentially explore regions of the state space that have higher probability.

Unlike the M-H model of movement, HMC makes use of local gradient information such that the walker (ant) tends to move in a direction of increasing probability. How *T. albipennis* may measure this is explored in the Discussion. For a system whose probability can be written in the form

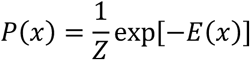

the gradient of *E*(*x*) can be evaluated and used to explore the probability space more efficiently. This is defined as:

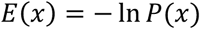

Using this definition the local gradient ∇*E*(*x*) can be calculated numerically.

The Hamiltonian is defined as *H*(***x, p***) = *E*(***x***) + *K*(***p***), where *K*(***p***) is a ‘kinetic energy’ which can be defined as:

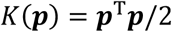

In HMC, this momentum variable ***p*** augments the state space ***x*** and there is an alternation between two types of proposal. The first proposal randomises the momentum variable, with ***x*** unchanged, and the second proposal changes both ***x*** and ***p*** using simulated Hamiltonian dynamics. The two proposals are used to create samples from the joint density

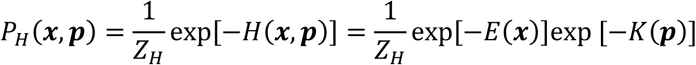

As shown, this is separable, so the marginal distribution of ***x*** is the desired distribution exp[−*E*(***x***)] /*Z*, and the momentum variables can be discarded and a sequence of samples {***x***^(*t*)^} is obtained that asymptotically comes from *P*(***x***) (Mackay, 2003).

We set the variable number of leapfrog steps (see supplementary Methods and Brooks et al., 2011)) to *L* = 10; after following the Hamiltonian dynamics for this number of steps a new momentum is randomly drawn and a new period of movement begins. This behaviour of moving intermittently in between updating the walker (ant) behaviour captures the behaviour observed in real ants (Hunt et al., 2016b) (see Discussion on gradient sensing). We set the leapfrog step length *ε* = 0.3 (see supplementary Methods for further introduction to L and *ε*).

For *N* = 18 simulated HMC ‘ants’ sampling from a sparse probability distribution (a gamma-distributed noise; see supplementary Methods), for 600 iterations, the r.m.s. displacement was again found and its log was regressed on log time. The gradient was found to be 0.567, 95% confidence interval (0.528 0.606), which is significantly greater than 0.5, so in this respect it is more similar to the superdiffusive search found in real ants (Franks et al., 2010).

We can also examine the correlation of velocities between successive movement periods. Since momentum ***p*** = *m****v*** is a vector in two-dimensional space, we can set *m* = 1 and find a magnitude for the momentum to determine the ‘speed’ of each movement (over the course of *L* = 10 leapfrog steps). In previous research on ant movements (Hunt et al., 2016b) the correlation between successive average event speeds in the cleaning treatment was found to be 0.407 ± 0.039 (95% CI). As expected for the HMC model, because the momentum is discarded and replaced with a new random momentum after each movement, the correlation of successive event speeds is equal to zero in this model. We can make the HMC model more ‘ant-like’ – and potentially more efficient – by only partially refreshing this momentum variable after the end of a movement period.

### Increasing correlations between steps: partial momentum refreshment (PMR)

HMC with one leapfrog step is referred to as Langevin Monte Carlo after the Langevin equation in physics (e.g. Kennedy, 1990) and was first proposed by Rossky et al. (1978). However, these methods do not require *L* = 1, so we use *L* = 10 to enhance comparability with the previous HMC model.

The momentum at the end of each movement can be updated according to the equation *p*′ = *αp* + (1 – *α*^2^)^1/2^*n*, where *p* is the existing momentum, *p*′ the new momentum, α is a constant in the interval [−1,1] and *n* is a standard normal random vector. With α less than one *p*’ is similar to *p* but with repeated iterations it becomes almost independent of the initial value. This technique of partial momentum refreshment (PMR) was introduced by Horowitz (1991). Such models are well-described in Brooks et al. (2011). Setting *α* equal to 0.65 (for example) and simulating with *N* = 18 results in speed correlations equal to 0.387 ± 0.012 (95% CI) which overlaps with the confidence interval for the real ant data.

The PMR method can be compared to an ant moving with a certain momentum (direction and speed) and then intermittently updating this momentum in response to its changing position in the physical and social environment, with a degree of randomness also included. The momentum changes as per the HMC method along a single trajectory, according to its subjective perception of foraging quality and potentially influenced by the pheromonal environment. If at the end of the trajectory it does not find itself in a more attractive region than before, it returns to its previous position: and with the correct model parameters (step size and number of leapfrog steps) this should be a relatively infrequent occurrence (see methodological discussion in Brooks et al. (2011)). Real ants have been predicted, and found, to leave ‘no entry’ markers when they turn back from an unprofitable location (Britton et al. 1998; Robinson et al. 2005). Its starting momentum in a particular direction is maintained to some degree but with some randomness mixed in – and so its previous tendency to move toward regions of high probability (quality) is not discarded as in HMC but used to make more informed choices about which direction to move in next. This is because foraging patches are likely to show some spatial correlation in their quality, with high quality regions more likely to neighbour other high quality regions (Klaassen et al., 2006; Van Gils, 2010). Previous empirical research (Hunt et al., 2016b) found evidence that ant movements are predetermined to some degree in respect of their duration. This implies that periods of movement are followed by a more considered sensory update and decision about where to move to next. A series of smaller movements (like 10 leapfrog steps) followed by a larger momentum update, as in the PMR model, would seem to correspond well with this intermittent movement behaviour.

### Measuring the performance of MCMC foraging models

The performance of the three MCMC models developed here can be measured in the following way. As discussed, the foraging ants should pursue a probability matching strategy, whereby they allocate their numbers across the environment in proportion to the probability that it will return (any) payoff. This will maximise the long-term rate of growth of the colony, or its biological fitness. Matching the probability distribution of resources in the environment can be understood as minimising the distance between it and the distribution of resource gatherers. In the domain of information theory, the difference between two probability distributions is measured using the cross-entropy

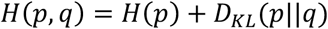

Where *H*(*p*) = − Σ_*i*_ *p*(*i*)log *p*(*i*) is the entropy of *p* and *D*_*KL*_(*p*‖*q*) is the Kullback-Leibler (K-L) divergence of q from p (also known as the relative entropy of *p* with respect to *q*). This is defined as

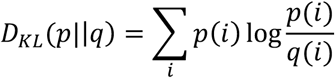

If we take *p* to be a fixed reference distribution (the probability of collecting resources in the environment), cross entropy and K-L divergence are identical up to an additive constant, *H*(*p*), and is minimised when *q* = *p*, where the K-L divergence is equal to zero. Cross-entropy minimisation is a common approach in optimization problems in engineering, and in the present case can be used to represent the task the ant foragers are trying to perform: match their distribution *q* with the distribution *p* of resources in the environment. The magnitude and rate of reduction of the cross-entropy is therefore used to compare the effectiveness of the MCMC models (M-H, HMC, PMR) presented here. However, as noted later, for dynamic environments (where the distribution of resource probabilities *p* is not fixed), K-L divergence is the suitable cost function to minimise.

Example simulations for the three models sample from a target distribution *p* with three simulated resources patches. This example distribution is generated by combining a gamma-distributed background noise (shape parameter=0.2, scale parameter=1) on a 100×100 grid given a Gaussian blur (*σ* = 3, filter size 100×100), what we refer to later as the ‘sparse distribution’, in equal 50% proportion with three patches of resources, which are single points of increasing relative magnitude of 1, 2, and 3 that have been given a Gaussian blur (*σ* = 10, filter size 100×100). The distribution *p* is thus also on a 100×100 grid. The simulations are run for 50,000 time steps, a reasonable period of time to explore this space of 10,000 points. Figure 2 shows the M-H model, which converges rather slowly on the target environment *p*. Figure 3 shows the performance of the HMC and PMR models, which show an improvement in the convergence rate because they avoid the random walk behaviour of the M-H model.

**Figure 2.**
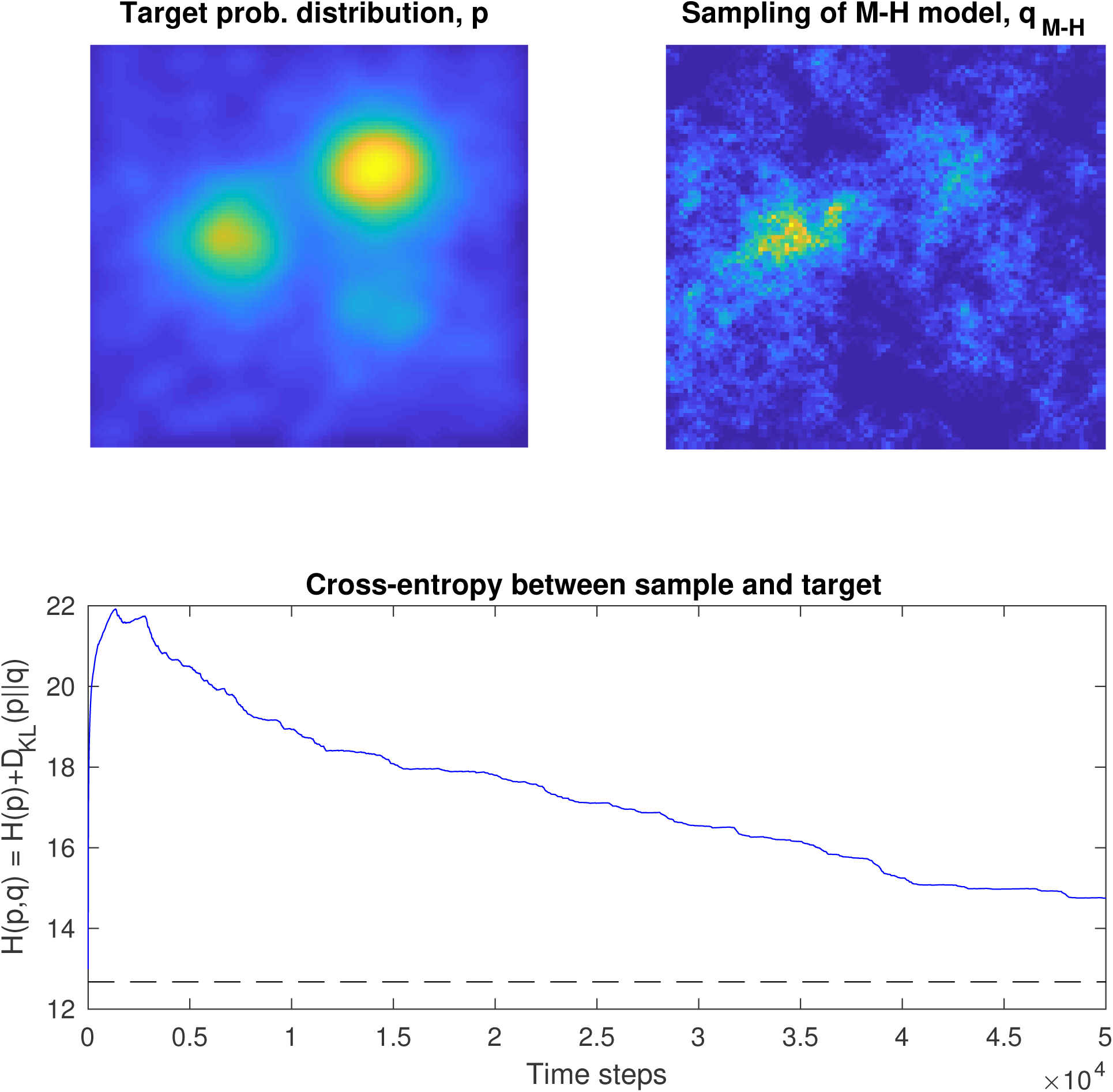
Performance of the M-H model as it generates a sample distribution *q* that approximates the target distribution *p*, the location of resources in the environment. The minimum cross entropy, where *q* = *p*, is shown as a dotted line.

**Figure 3.**
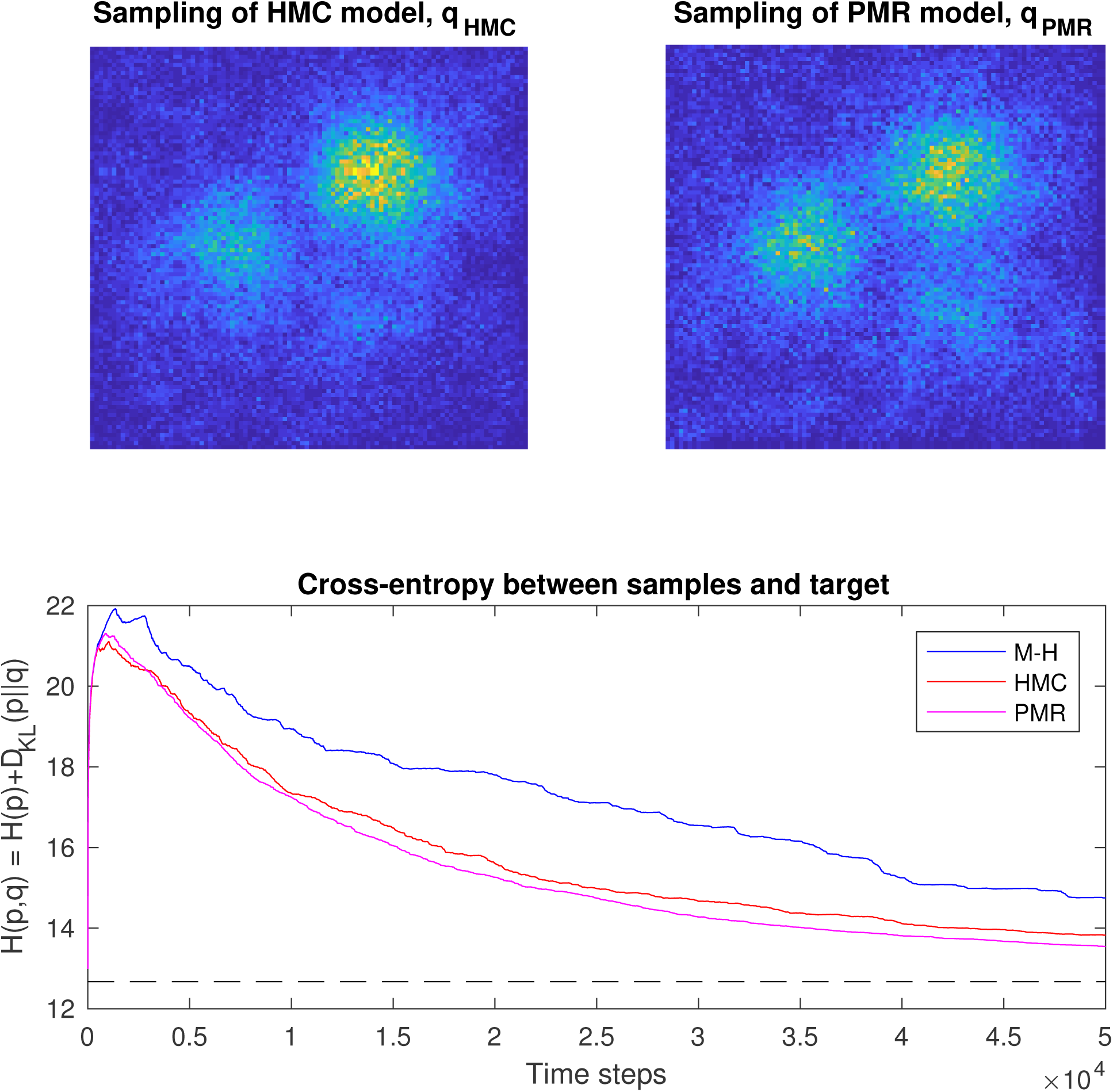
Performance of the HMC and PMR models, compared to that of M-H. In general, HMC and PMR outperform M-H because random walk type exploration of probability space is avoided, by following local gradient information and making larger steps. Their performance depends on the nature of the target distribution and choosing suitable values for step length *ε* and number of steps *L*.

Figure 4 shows example trajectories from real ants (Hunt et al. 2016a) for a period of 100 s, and for 100 timesteps of the 3 models. The ants are in an empty arena and the models are sampling from a sparse distribution (supplementary Methods). The random walk behaviour of the M-H model is evident, while the greater tendency to make longer steps in one direction is evident in the PMR model in comparison to the HMC model.

**Figure 4.**
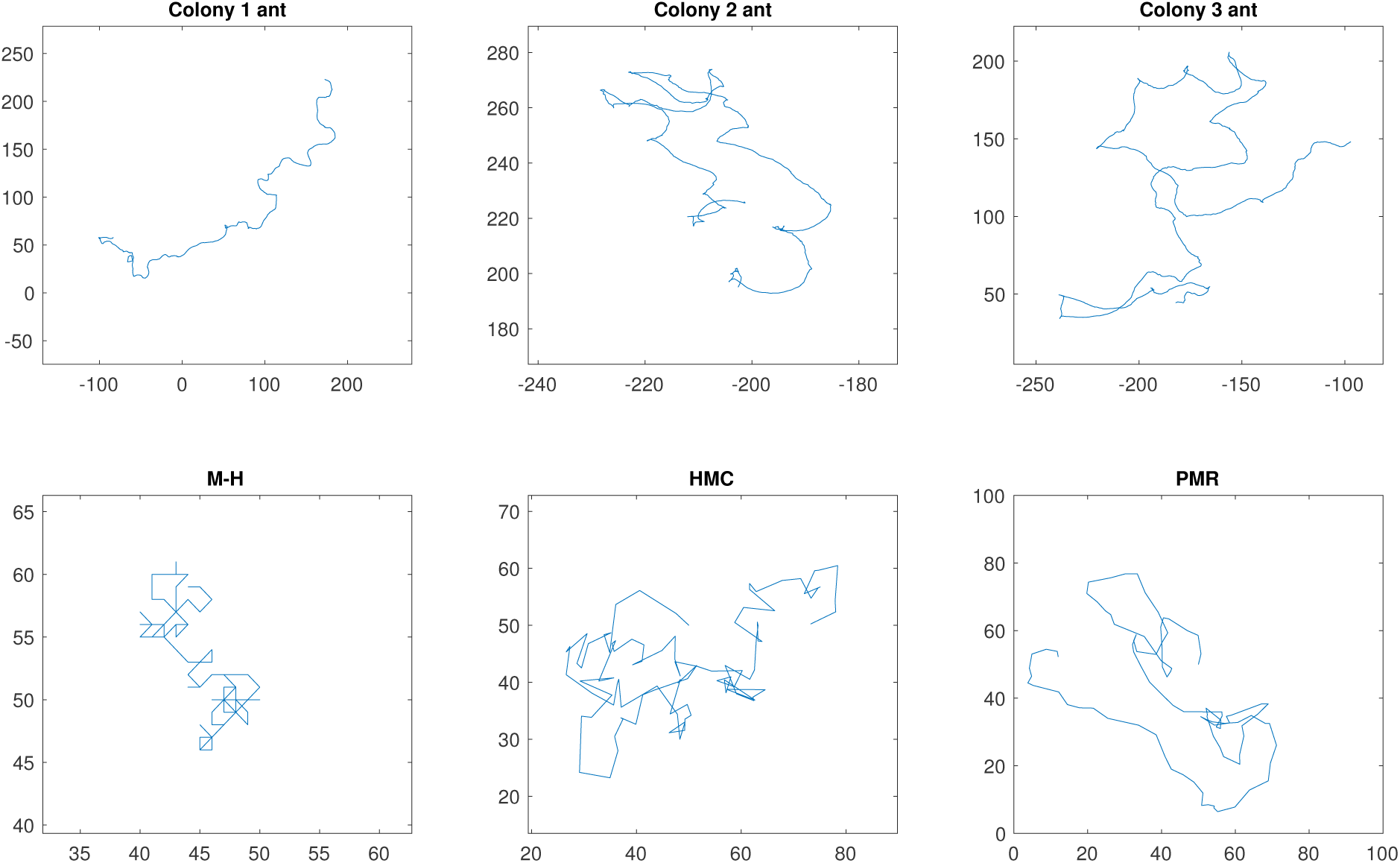
Comparison of example trajectories from real ants and for the 3 MCMC models (100 simulated timesteps). The model trajectories become increasingly superdiffusive.

Figure 5 shows the distribution of directional changes (change in angle heading) between steps. The distribution of direction changes is known as the phase function in statistical physics and has been applied to ant trajectory analysis by for instance Khuong et al. (Khuong et al. 2013). Real ants can make large changes of direction, of course, but this is rarely done with an abrupt heading shift. The M-H model moves grid-wise in single steps; the HMC model has no correlation between step directions; while the PMR model tends to make each new step in a similar (correlated) direction to the prior one. In this respect, too, PMR is a better model of ant movement.

**Figure 5.**
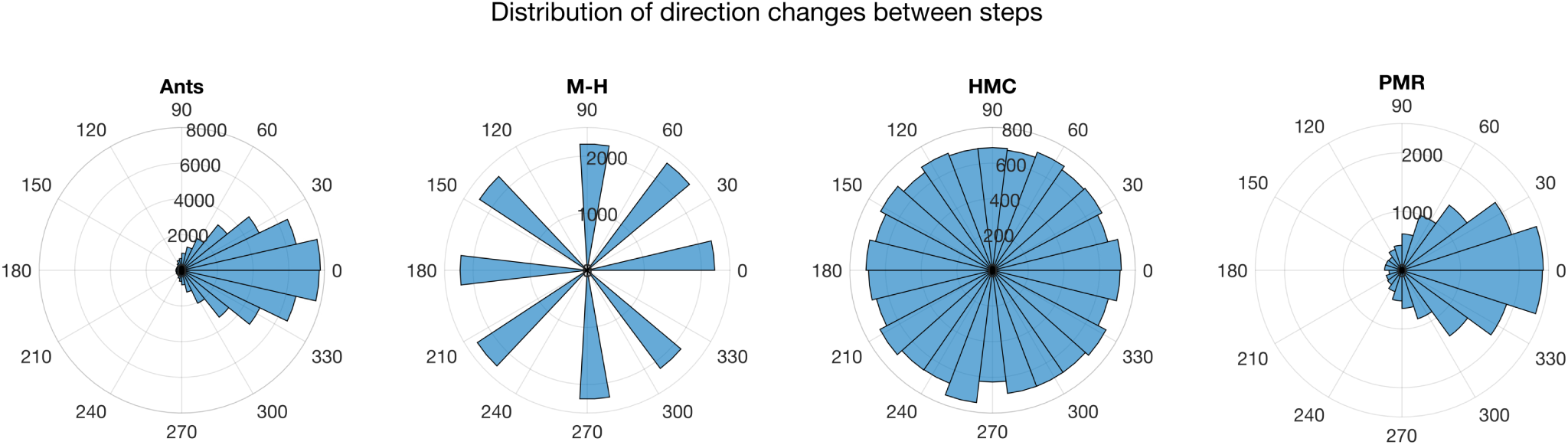
The distribution of direction changes between steps in real ants (N=18) and the three MCMC models (simulated for N=18 ‘ants’ for 1000 timesteps).

### Optimal foraging and Lévy flights

We have presented a new class of foraging models based on MCMC methods, which operationalise movement for a Kelly strategy (probability matching) in two-dimensional space. There is an extensive theoretical and empirical literature examining the distribution of step lengths for foraging animals that considers the hypothesis that a Lévy distribution is optimal (Bartumeus, 2007; Benhamou, 2007; Humphries et al., 2010; Viswanathan et al., 2008). Lévy flights are a particular form of superdiffusive random walk where the distribution of move step-lengths fits an inverse power law such that the probability of a move of length *l* is distributed like (*l*) ≈ *l*^−*μ*^, where 1 < *μ* ≤ 3.

We use the method of Humphries et al. (2013) to identify individual movement steps in two dimensional data, treating monotonic movements in a certain direction in one dimension (i.e., *x* or *y*) as a step. We estimate the exponent using maximum likelihood estimation (White et al., 2008). The distribution of ranked step length sizes in both real and simulated data is shown in Figure 6.

**Figure 6.**
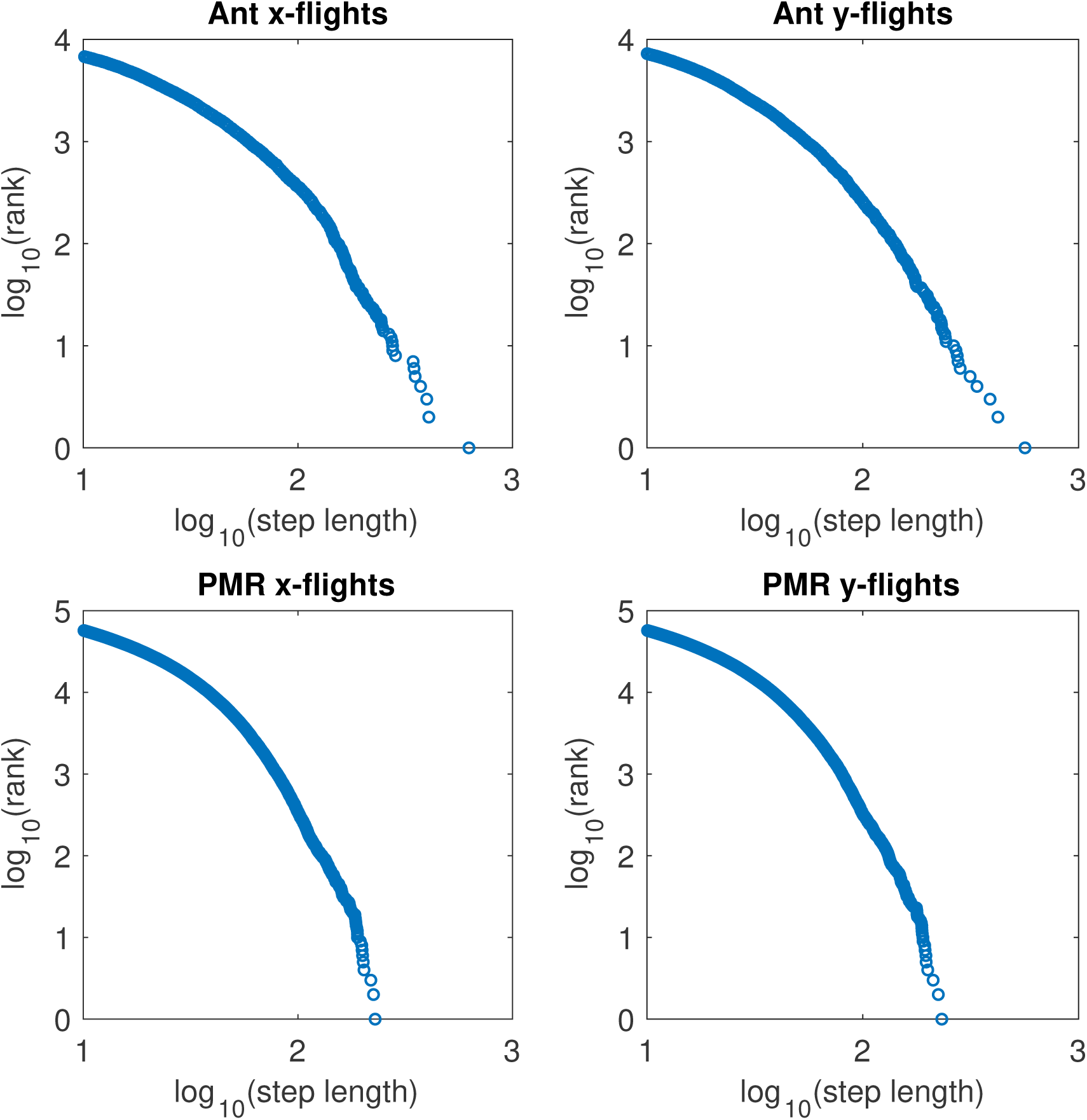
The (apparently) power-law distributed step lengths for both real ants and simulated PMR walkers.

There are similar exponents estimated (Table 1) for both the real ant data in an empty arena (N=18 ants from 3 colonies) and PMR trajectories (100 ‘ants’ for 5000 iterations) sampling from a sparse probability distribution (supplementary Methods). The exponent *μ* in both cases is in the right region for a Lévy flight 1 < *μ* ≤ 3. This would seem to be evidence for a Lévy strategy in the ants (though variation in individual walking behavior can also contribute to the impression of a Lévy flight (Petrovskii et al. 2011)), but we suggest an alternative in the next section of this paper.

**Table 1.** Power-law exponents in both empirical and PMR simulated trajectories potentially indicate a Lévy walk

|  |  |  |  |
| --- | --- | --- | --- |
| Dimension of steps | Maximum likelihood estimate of exponent, truncated Pareto distribution |  | 402 |
|  | Empirical data | PMR data | 403 |
| $x$ | 2.41 | 2.26 | 404 |
| $y$ | 2.55 | 2.26 | 405 |

## Discussion

### MCMC models and existing movement paradigms

The framework we develop here is an important step in integrating key perspectives in movement research, as described for example by Nathan et al. (2008). It incorporates elements of randomness, producing correlated random walks in certain environments; it quantifies optimality in respect of foraging strategies via cross-entropy (Kullback-Leibler divergence); it includes an important aspect of common animal behaviour, namely intermittent movement (Kramer and McLaughlin, 2001), and specifically for the ants’ neural and/or physiological behaviour, apparent motor planning (Hunt et al., 2016a); and it makes explicit the information used by the animal step-by-step. Finally, and crucially, it explicates cognition at the emergent group level, because individual movement is at the service of a group-level Kelly strategy. One component of Nathan et al.’s framework is the internal state of the organism. This is not included in the models here, though state-dependent behaviours such as tandem running (Franks and Richardson, 2006) could be included by analogy with particle filtering (e.g. Gordon et al., 1993), for instance (Hunt et al., in prep, 2018b). Our use of the Markov assumption (movement being memoryless, depending only on the current position) is justifiable with respect both to the worker ant’s individual cognitive capacity, and its single-minded focus on serving the colony through discovering and exploiting resources. Its motion capacity is linked to the specification of a partial momentum refreshment model; while we specify the navigation capacity in its ability to measure the quality gradient, which is also an externally determined factor.

### The mechanisms and challenges of gradient sensing

We may consider further the ability of ants to use local gradient information, as in the HMC and PMR models, with respect to the ants’ sensory system. *Temnothorax albipennis* is well-known for relying heavily on visual information in movement (McLeman et al., 2002; Pratt et al., 2001) and in common with most (or perhaps all) ants on olfactory information. It may be that the intermittent movement examined in Hunt, Baddeley et al (Hunt et al., 2016b) is associated with limitations in the quality of sensory information when moving (Kramer and McLaughlin, 2001). We suggest that *T. albipennis* workers have relatively good eyesight for a pedestrian insect and their small size, having around 80 ommatidia in each compound eye (Hunt et al., 2018a) and may be assumed conservatively to have an angle of acuity of 7 degrees (Pratt et al., 2001). Therefore, movement would seem unlikely to make much difference to how well they can see. Since our model highlights the importance of gradient following, this may be more difficult to measure for the olfactory system during movement. Indeed, in Hunt et al. (2016b) we suggest that social information from pheromones or other cues is only fully attended to during periods of stopping because of motor planning, with the duration of movements being predetermined by some endogenous neural and/or physiological mechanism. This may be therefore a mechanistic reason for the stepwise movement in the PMR model, in addition to its informational efficiency which is its evolutionary origin. Even more sophisticated MCMC models that rely on the second derivative of the probability distribution, such as the Riemann Manifold Langevin method (Girolami and Calderhead, 2011), may be relevant, because this property (the rate of change of the gradient) may be only measured with adequate accuracy when the ant is at rest.

### Lévy step length distributions indicate a world with little gradient information

In a ‘flat’ quality landscape, or sparse world, our model generates Lévy-like behaviour as seen for instance in Reynolds et al., 2013. This remains an adaptive response, but it is not a true Lévy distribution, because there is a finite variance. Much interest has been generated by Lévy flight based foraging models which theoretically optimise mean resource collection for certain random worlds; and this would seem to be evidence for just such a strategy in *T. albipennis* ants. Yet here we make a simple point that rather than being a deliberate strategy, Lévy-like behaviour may result from an organism lacking cues about which way to move. Scale-free reorientation mechanisms have indeed been suggested as a response to uncertainty in invertebrates (Bartumeus and Levin, 2008). Yet the generation of a Lévy-like distribution from our gradient-following model suggests that such observations may not really be scale-free. The empirical distribution of momentums provides insight into the length-scales on which the world remains smooth.

### Measuring efficiency, selecting for unpredictability

The rate of resource collection can be straightforwardly calculated by finding the cross-entropy (Kullback-Leibler divergence) between the spatial distribution of resources, and the realised foraging distribution resulting from the foraging strategy. The distribution of resources is seen from a ‘genes-eye’ view of the animal or superorganism, with respect to maximising the long-term biomass or number of copies of genes in the environment: this focuses on a location’s probability of yielding resources, or reliability, as opposed to the one-off payoff. The foraging strategy is that chosen by natural selection. Minimising the cross-entropy (Kullback-Leibler divergence) is achieved by obeying the matching law: foraging proportional to the probability finding the best resources at each available location. This strategy is especially suited to a superorganism like an ant colony, because it can forage in multiple locations simultaneously by allocating worker ants in numbers proportional to the location’s reliability, through self-organisation (Camazine et al., 2001).

There has been some intimation before that MCMC could be a model for biological processes (Neal, 1993), with some query about whether the requisite randomness is possible in organisms. We think that not only is spontaneous (i.e. non-deterministic, ‘random’) behaviour present, it is necessary for survival in terms of being unpredictable around predators, prey or competitors (Brembs, 2010), or for finding food using a ‘strategy of errors’ (Deneubourg et al., 1983). Indeed, the Bayesian framework developed here allows predictions to be developed regarding the optimal amount of ‘randomness’ in behaviour at both the level of the individual and colony (in the partial momentum refreshment model, this is adjusted with the *α* parameter) that can be tested in future empirical research. Further predictions arise from the momentum reversal step in MCMC PMR (Brooks et al., 2011), which may be compared to observations of U-turning in ants (Beckers et al., 1992; Hart and Jackson, 2006). Recent literature (Hoffman and Gelman, 2014) has developed methods to adjust the path length dynamically, while removing the need to have a parameter *L* for the number of leapfrog steps. Observing how ants (and other organisms) adjust their step lengths according to different resource distributions will be instructive of their underlying movement model.

### Selection for collective foraging phenotypes

The major evolutionary transitions (Maynard Smith and Szathmary, 1995) can be seen as successive leaps forward in information processing efficiency. The Bayesian framework developed here permits the evaluation and prediction of alternative movement strategies, for groups of high-related organisms, in quantitative, informational terms, in relation to environmental resource distributions. Our framework permits us to make the simple statement that for a movement strategy to be favoured under natural selection:

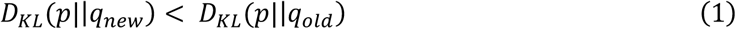

i.e. the Kullback-Leibler divergence (measuring the similarity of two distributions) between a potential (genetically accessible) collective movement strategy that results in the equilibrium distribution of foragers *q*_*new*_, and the organism’s resource environment *p*, has to be lower than under the current strategy found in the population that results in distribution *q*_*old*_. This reduction may indeed be achieved by more sophisticated, coordinated, collective behaviour, notwithstanding higher individual energetic cost. Future research could relate such an expression to concepts in evolutionary genetics such as fitness landscapes (Orr, 2009). The theoretical relationship between the level of relatedness within a social group, and the relevance of the Kelly strategy, could be explored in future research.

## Conclusion

We described the foraging problem as a repeated multiplicative game, where an ant colony has to place ‘bets’ on which foraging patches to visit, with an ultimate payoff of more colonies or copies of their genes being created. Ants are very successful in terms of their terrestrial biomass (Schultz, 2000), and so it would seem likely that they are following a highly evolved strategy. We suggest the theoretical optimum is a ‘Kelly’ or probability matching strategy, which maximises the long term ‘wealth’ or biomass of the colony rather than the resource collection of single ants. By mapping the foraging problem to a set of methods designed to effectively sample from probability distributions, we present models of ant movement that achieve this matching behaviour. These MCMC-based models thus provide spatially explicit predictions for movement that describe and explain how colonies optimally explore and exploit their environment for food resources. We also show how Lévy-like step length distributions can be generated by following a local gradient that is uninformative, suggesting that contrary to being an evolved strategy, Lévy flight behaviour may be a spontaneous phenomenon. While we do not include interactions between ants in the model, past theoretical analysis of information use in collective foraging suggests that totally independent foraging is actually optimal for a broad range of model parameters when the environment is dynamic. This is because information about food patches may not be worth waiting for if they are short lived (Dechaume-Moncharmont et al., 2005).

Understanding the logic of information flow at the level of the gene and the cell has been identified as a priority (Nurse, 2008). However, given that no level of organisation is causally privileged in biology (Noble, 2011), explicating this at the organismal and super-organismal level should also advance our understanding. Our Bayesian framework operationalizes earlier proposed frameworks (such as that of Nathan et al., 2008) for movement in a coherent and logical way, accounting for the uncertainty in both the individual ant and the colony’s cognition in relation to the foraging problem. It also allows quantification of the system’s emergent information processing capabilities and hypothesis generation for different organisms moving in different environments. Our MCMC models can be used as a foundation upon which further organismal and ecological complexity can be explained in future research; and suggest that the movement strategies of animal collectives may be instructive for biomimetic improvements to MCMC methods.

## Author contributions

ERH drafted the paper, produced the analysis and simulations, and suggested eqn. (1); NRF advised on social insect biology; RJB proposed the Bayesian framework and its specific theoretical concepts; all authors contributed to the present draft.

## Funding statement

E.R.H. thanks the UK Engineering and Physical Sciences Research Council (grants no. EP/I013717/1 to the Bristol Centre for Complexity Sciences, EP/N509619/1 DTP Doctoral Prize).

## Ethics statement

The ant *Temnothorax albipennis* is not subject to any licencing regime for use in experiments. The ants were humanely cared for throughout the experiment.

## Competing interests

We have no competing interests.

