## Supplementary Methods for "Optimal foraging and the information theory of gambling"

***Further details on Kelly betting***

For full treatment see chapter 6 of Cover and Thomas (2006). A probability matching strategy is also optimal with 'superfair' odds when $\sum \frac{1}{\sigma_i} < 1$. In both of these cases, the full capital or proportion of foragers should be allocated across foraging sites to maximise the long-term wealth, although the Kelly strategy may also be followed by betting a fixed proportion $\alpha$ of full bets. This so-called $\alpha$-Kelly strategy has a much slower rate of growth in wealth (MacLean et al., 2010), however, short-term volatility in wealth (or biomass) is reduced by a factor $\alpha^2$ (Rising et al., 2012). Animals are often risk averse to variable rewards (Kacelnik and Bateson, 1996) and this may be seen at the level of the colony; in practice only a proportion of ants forage, with some staying purposefully inactive for significant periods of time (Charbonneau and Dornhaus, 2015). In a colony of, say, 200 workers, 50 might forage, which would correspond to $\alpha = 0.25$; but for the purpose of this paper we do not consider this further.

When $\sum \frac{1}{\sigma_i} > 1$ subfair odds are said to be available, and it is better to hold back some proportion of the ant foragers (or a gambler's wealth) and try to bet on outcomes offering the most favourable expected return $p_i \sigma_i$. In this case the optimal betting amount is solved using a 'waterfilling' method and is dealt with in Erkip (1996).

***MCMC methods – resources***

Good introductions to Markov chain Monte Carlo (MCMC) methods can be found in Neal (1993), Mackay (2003), and Brooks et al. (2011).

***Metropolis-Hastings – further details and comparison with ant data***

The M-H method makes use of a proposal density $Q$ (which depends on the current state $x$) to create a new proposal state to potentially sample from. Q can be simply a uniform distribution: in a discretized environment these can be $x^{(t)} + [-1,0,1]$ with equal probability. To decide whether to accept the new state as the next step in the sample, we compute the quantity

$$a = \frac{P^*(x')}{P^*(x^{(t)})} \frac{Q(x^{(t)}; x')}{Q(x'; x^{(t)})}$$

If $a \geq 1$ then the new state is accepted; otherwise it is accepted with probability $a$. If the step is accepted, we set $x^{(t+1)} = x'$ ; if it is rejected then $x^{(t+1)} = x^{(t)}$ (the current state is taken forward to the next step). The M-H method is widely used for sampling from high-dimensional problems, but it has a major disadvantage that it explores the probability distribution by a random walk, which can take many steps to move through the space, according to $\sqrt{T}\epsilon$ where T is the number of steps and $\epsilon$ is the step length.

The Metropolis-Hastings algorithm was used to sample from a sparse probability distribution. This is generated by combining a gamma-distributed background noise (shape parameter=0.2, scale parameter=1) on a 100×100 grid given a Gaussian blur ($\sigma = 3$, filter size 100×100) Using $N = 18$ simulated walkers exploring for 600 iterations, the exponent was found to be 0.487, 95% confidence interval (0.444 0.529), i.e. not significantly different to 0.5 and so engaging in a standard diffusive search, as expected. We can use the M-H walkers as a model of exploring ants, and so to assess its validity we can examine trajectory data of real ants exploring a uniform empty space (Hunt et al., 2016a) and compare it with simulations from this model, to see whether ants are engaged in a standard diffusive (Brownian) exploration of space. The root mean square (r.m.s.) displacement for $N = 18$ ants was found for the first 60 seconds of their exploration of an unfamiliar arena (600 trajectory points with a sampling period of 0.1s, a time short enough that they would not reach the edge of the arena), and its log was regressed on log time. The gradient was found to be 0.59 with a 95% confidence interval of (0.55,0.63), which is greater than 0.5 and so the ants are engaged in superdiffusive search.

***Leapfrog method***

In two of the MCMC methods presented, HMC and PMR, there are two key parameters ε and $L$ that must be chosen suitably to generate a sample efficiently. See Brooks et al. (2011) for a full treatment of these issues.

Although the dynamics of a system in Hamiltonian mechanics is described using differential equations for continuous time and space, for implementation on a computer the equations must be approximated by discretizing time using a small step length ε. We may assume a Hamiltonian of the form $H(q,p) = U(q) + K(p)$ and a kinetic energy $K(p) = p^T M^{-1} p/2$, and a diagonal M with elements $m_1, \ldots, m_d$ so that $K(p) = \sum_{i=1}^{d} \frac{p_i^2}{2m_i}$ . Then the leapfrog method proceeds as follows:

$$p_i\left(t + \frac{\varepsilon}{2}\right) = p_i(t) - (\varepsilon/2)\frac{\partial U}{\partial q_i}\big(q(t)\big),$$

$$q_i(t+\varepsilon) = q_i(t) + \varepsilon\frac{p_i(t+\varepsilon/2)}{m_i},$$

$$p_i(t+\varepsilon) = p_i(t+\varepsilon/2) - (\varepsilon/2)\frac{\partial U}{\partial q_i}\big(q(t+\varepsilon)\big).$$

We start with a half step for the momentum variables, a full step follows for the position variables using those new momentum variables, and finally we do another half step for the momentum variables using the new values for the position variables. This process is iterated a certain number of times $L$, and preserves volume exactly. It is also reversible by negating $p$ and applying the same number of steps $L$, then negating p again. It gains its name from the use of interleaved time points such that $p_i$ and $q_i$'leapfrog' over each other.

The leapfrog step length was set to be $\varepsilon = 0.3$ in the simulations presented in the main text. Selecting a suitable step length is vital, because if it is too large there will be a very low acceptance rate for the states proposed by the simulated trajectories. If it is too small it will waste computation time, as the trajectories will make unnecessarily short progress through the space, or it can lead to slow exploration by a random walk through the space if the trajectory length $\varepsilon L$ is too small, which defeats the object of the method. In the first instance reasonably effective ε and $L$ can be found through trial and error.
